# Dynamic and topological properties of large-scale brain networks in rapid eye movement behavior disorder

**DOI:** 10.1101/2024.05.25.595917

**Authors:** Yidi Li, Kenji Yoshinaga, Takashi Hanakawa, Japan Parkinson’s Progression Markers Initiative (J-PPMI) study group

## Abstract

**Introduction:** There is a lack of research in the existing literature when it comes to analyzing the dynamics of resting-state functional magnetic resonance imaging to understand the underlying mechanisms of isolated rapid eye movement sleep behavior disorder (iRBD). This study aims to contribute to our understanding of abnormalities in brain network dynamics in iRBD and their association with alpha-synucleinopathy. Additionally, I employed graph theoretical metrics to obtain a topological insight into the brain network of iRBD.

**Methods:** Resting-state fMRI data from 55 iRBD patients and 97 healthy controls (HCs) were utilized. A sliding window approach, functional connectivity analysis, and graph theory analysis were applied to the data. I calculated the mean, standard deviation, skewness, and kurtosis of the time series for both dynamic functional connectivity (dFC) and four graph metrics (clustering coefficient, global efficiency, assortativity coefficients, and eigenvector centrality). Subsequently, I compared the those metrices between iRBDs and HCs. Relationships between clinical scales and abnormal dFC were assessed using a general linear model.

**Results:** iRBD patients exhibited abnormal mean dFC, particularly in the default mode network, sensorimotor network, basal ganglia network, and cerebellum. Kurtosis of dFC revealed abnormalities between the middle temporal gyrus and cerebellum. Group differences were also observed in the mean eigenvector centrality of the precentral gyrus and thalamus.

**Conclusion:** The mean of dFC identified impairments putatively in movement functions and various compensatory mechanisms. Moreover, mean eigenvector centrality revealed topological changes in motor-related network in iRBDs. The use of kurtosis as a potential index for extracting dynamic information may provide additional insights into pathophysiology in iRBDs.

## 1. Introduction

Rapid eye movement (REM) sleep behavior disorder (RBD) is characterized by abnormal REM sleep without muscle atonia, and it is termed isolated RBD (iRBD) when occurring independently of other diseases. Multiple lines of evidence^1–3^ suggest iRBD as a prodromal form of alpha-synucleinopathy, emphasizing the need to identify early neuropathological changes of iRBD and its relationship to neurodegenerative diseases. Thus, individuals with iRBD face a significant risk of progression to conditions including dementia with Lewy bodies, multiple system atrophy, and Parkinson’s disease (PD).

Resting-state functional magnetic resonance imaging (rs-fMRI) is widely used to search for pathophysiological correlates of neurological and psychotic disorders. Commonly applied methods to resting-state data assume static interactions between regions during the scan duration. Using various static analysis methods including functional connectivity (FC) analysis, amplitude of low frequency fluctuations (ALFF) analysis, regional homogeneity (ReHo) analysis and so on, rs-fMRI studies have found abnormality related to the function of the motor cortex^4,5^, basal ganglia^6,7^, occipital lobe^7,8^, prefrontal cortex^9^ and cerebellum^10,11^. Importantly, rs-fMRI results also proven to related to clinical and neuropsychological data. Thalamo-fusiform FC value was found to positively correlated with word list recognition^7^, decreased FC of the central autonomic network were negatively correlated with the Scales for Outcomes in Parkinson’s Disease–Autonomic scores^11^. In a study using graph theory analysis, nodal efficiency in the bilateral thalamus were positively correlated with RBD screening questionnaire(RBDSQ) scores^12^.

While contributing significantly to understand physiopathology of iRBD and its relationship to alpha-synucleinopathy, these static approaches face criticisms because they neglect temporal fluctuations of rs-fMRI time-series^13^. Intrinsic fluctuations in brain activity have long been acknowledged, with even greater prominence during the resting state when mental activity is unconstrained^14^. Though a pioneer study has applied dynamic analysis methods to iRBD patients’ rs-fMRI data and identified different temporal characteristic of brain states compared to healthy controls^15^, there is still a lack of studies and consensus on the dynamics of resting-state networks in iRBDs, encompassing the methods used to extract temporal fluctuations, the reproduction and consistency of results. Therefore, in this study, I calculated dynamic connectivity and graph-theoretical metrics to fill the gaps in the dynamic aspect of rs-fMRI data analysis in iRBD. This study is a part of Japan Parkinson’s Progressive Marker Initiative (J-PPMI)^16^.

## 2. Materials and Methods

### 2.1 Participants

We analyzed cross-sectional MRI, clinical and neuropsychological data acquired by the National Center of Neurology and Psychiatry (NCNP) site of in the J-PPM multi-site cohort. The dataset was available in an anonymous format, including 55 individuals diagnosed as iRBD (mean age: 69.7 ± 5.8 years, age range: 60-83 years; 39 males, 16 females) and 97 healthy controls (HCs) (mean age: 66.5 ± 8.7 years, age range: 41-83 years; 47 males, 50 females). The diagnosis of iRBD adhered to the criteria outlined in the International Classification of Sleep Disorders, third edition (ICSD-3)^17^. All individuals with iRBD underwent overnight polysomnography, confirming REM sleep without atonia.

All participants gave written informed consent to participate in the study approved by the NCNP ethics committee (A2014-127 and A2018-056) following the declaration of Helsinki. The present study is a reanalysis of the data set used for machine learning-based classification between HC and iRBD (Matsushima et al. 2024).

### 2.2 Data acquisition

#### 2.2.1 MRI data acquisition

MRI data were acquired using a 3-T MR scanner with a 32-channel phased array head coil (MAGNETOM Verio Dot, Siemens Medical Systems, Erlangen, Germany). Resting-state fMRI data were obtained using gradient-echo echo planar imaging (GE-EPI) for 10 min, during which participants were asked to clear their minds and focus on a central fixation cross. To make sure that participants remained awake during the resting-state fMRI scanning, the Stanford sleepiness scale^18^ was administered. The acquisition parameters for GE-EPI were as follows: repetition time (TR) = 2500 ms, echo time (TE) = 30 ms, flip angle (FA) = 80°, voxel size = 3.3 × 3.3 × 3.2 mm^3^ (with a 0.8-mm gap), 40 axial slices, and a total of 240 volumes. A double echo gradient echo sequence was used to provide an MRI field-map in the same space with GE-EPI as follows: TR = 488 ms, TE1 = 4.92 ms, TE2 = 7.38 ms, FA = 60°, voxel size = 3.3 × 3.3 × 3.2 mm^3^ (with a 0.8-mm gap), and 40 axial slices. Additionally, structural MRI data were acquired using a three-dimensional T1-weighted magnetization-prepared rapid gradient-echo (MPRAGE) sequence with the following parameters: TR = 1900 ms, TE = 2.52 ms, inversion time (TI) = 900 ms, FA = 90°, 192 sagittal slices, and voxel size = 0.98 × 0.98 × 1 mm^3^.

#### 2.2.2 Clinical and neuropsychological assessments

In people with iRBDs, RBD episodes and daytime sleepiness in daily life were screened using the Japanese edition of RBD screening questionnaire (RBDSQ-J)^19^ and the Epworth Sleepiness Scale^20^, respectively. The Movement Disorder Society-sponsored revision of the unified Parkinson’s disease rating scale (MDS-UPDRS) part III score was recorded for individuals with iRBD. A MDS-UPDRS part III score below 7 (excluding the kinetic tremor and postural tremor score) was taken as evidence arguing against parkinsonism^21^. All iRBDs and 18 HCs underwent the Japanese version of the Montreal cognitive assessment (MoCA-J)^22^ as a screening test for cognitive functions. All HC were evaluated using the mini-mental state examination (MMSE).

### 2.3 Data analysis

#### 2.3.1 Preprocessing

The preprocessing of structural MRI and functional MRI data was carried out primarily using the CONN toolbox (https://web.conn-toolbox.org/). The preprocessing pipeline included realignment and unwarping, slice-timing correction, outlier identification, tissue segmentation, spatial normalization, and smoothing (6 mm full width at half maximum)^23^. Denoising was performed using the anatomical component-based noise correction procedure (aCompCor) followed by FMRIB’s ICA-based Xnoiseifier (FIX), ensuring vigorous removal of noise components.

Volumes of interests (VOIs) were defined for 91 cortical areas and 15 subcortical areas from the FSL Harvard-Oxford Atlas, as well as 26 cerebellar areas from the AAL atlas.

#### 2.3.2. Dynamic analysis

##### 2.3.2.1. Sliding window analysis

Sliding window analysis was conducted using the sliding window approach facilitated by the DynamicBC toolbox (http://restfmri.net/forum/DynamicBC). The sliding window approach is a widely adopted strategy for quantifying the time-varying behavior of the chosen metric throughout the scan. With this method, considering a scan with N time points, a temporal window of fixed length W was selected. Data points within this window, ranging from time t=1 to t=W, were utilized to calculate matrices of interest. Subsequently, the window was shifted in time by a fixed number of data points, referred to as steps (T), and the same calculation was conducted over the time interval [1+T, W+T]. This process was repeated until the window spanned the entire time points, resulting in S=((N-W)/T) +1 windows^24^. In this study, we chose rectangular window sizes—20 TRs (50 seconds)—with a step size of 1 TR (2.5 seconds), resulting in a total of 221 windows.

##### 2.3.2.2. Dynamic functional connectivity analysis

A functional connectivity (FC) matrix was established by calculating Pearson’s correlation coefficients between the time series of each of the 132 VOIs across all 152 subjects.

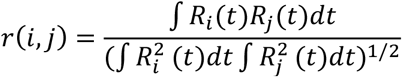

Where *R* is the rs-fMRI timeseries within each VOI, *r* is a FC matrix, and *i,j* are indices of VOIs. Dynamic functional connectivity (dFC) was computed for each of the 221 windows (window size of 20 TRs), producing 221 dFC matrixes in each of the 152 subjects using DynamicBC toolbox (http://restfmri.net/forum/DynamicBC).

We then computed four summary statistics (mean, standard variance, skewness, and kurtosis) for each dFC time series (132*132*221), to providing temporal information.

Skewness quantifies the asymmetry of the data relative to the sample mean. A negative skewness indicates that the data is more spread out to the left of the mean than to the right. Conversely, a positive skewness implies that the data is more spread out to the right. For a perfectly symmetric distribution, such as the normal distribution, the skewness is zero. The skewness of a distribution is defined as:

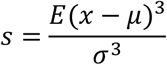

where *µ* is the mean of *x, σ* is the standard deviation of *x*, and *E(x)* represents the expected value of the quantity *t*. (https://ww2.mathworks.cn/help/stats/skewness.html)

Kurtosis serves as a measure of the frequency of outlier occurrences. The kurtosis of a normal distribution is 3. Distributions with kurtosis greater than 3 are more prone to outliers than the normal distribution, while those with kurtosis less than 3 are less susceptible to outliers.

Similarly, the kurtosis of a distribution is defined as:

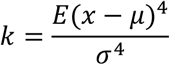

where *μ* is the mean of *x, σ* is the standard deviation of *x*, and *E(x)* represents the expected value of the quantity *t*. (https://ww2.mathworks.cn/help/stats/kurtosis.html)

##### 2.3.2.3. Graph theorical analysis

A graph comprises of nodes and edges. In the context of resting-state networks, we defined nodes by specific brain areas (VOIs in this study) and edges by the functional interactions across a pair of these brain areas (dFC in this study).

Before passing the dFC matrixes to the graph theorical analysis, the dFC matrixes were thresholded using significant correlation coefficients (P<0.05, FDR corrected) to eliminate spurious connections and generate sparse matrices for subsequent topological analysis.

As topological metrices, we calculated clustering coefficient (C), global efficiency (GE), assortativity coefficients and eigenvector centrality (EC) with dFC matrices using the Brain Connectivity Toolbox (http://www.brain-connectivity-toolbox.net) in each participant.

The clustering coefficient (C) is defined as the fraction of a node’s neighbors that are also neighbors of each other^25^ and is a measure of how well-connected and interrelated the immediate neighbors of a node are. Mathematically, C is expressed as follows^26^:

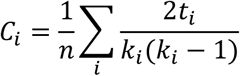

where *n* is the number of nodes in the graph, *t*_*i*_ is the number of triangles around a node *i* and *k*_*i*_ means the degree of a node *i*. The C for the network therefore characterizes the prevalence of clustered connectivity around individual nodes.

The global efficiency (GE) is a measure for overall integration and communication efficiency of a network and reflects how well information can be transmitted and integrated across all pairs of nodes in a network^26^. GE is defined as:

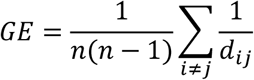

where *d*_*ij*_ represents the shortest path between the nodes *i* and *j*. GE can be meaningfully computed even in disconnected networks where paths between disconnected nodes are considered to have infinite length, leading to zero efficiency^26^.

The assortativity coefficient helps characterize the organization and resilience of network and measures how anatomical features influence network vulnerability to insults. It gauges the extent to which nodes in a network associate with other nodes of similar or opposing types^27^. Mathematically, assortativity coefficient is defined as the correlation coefficient between the degrees of all nodes on two opposite ends of a link:

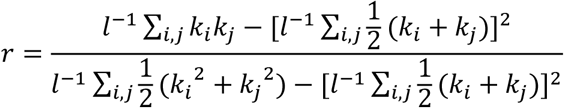

where *l* is a number of the links in the network. Networks with a positive assortativity coefficient are more likely to have a resilient core of mutually interconnected high-degree hubs. Conversely, networks with a negative assortativity coefficient are likely to possess widely distributed and consequently vulnerable high-degree hubs^26^.

The eigenvector centrality (EC) is one of the measures for node centrality and indicates the importance of nodes based on their connections with other important nodes in the network. It evaluates the centrality of a node not only by the number of its connections but also by the importance of those connections insomuch as to take the whole network into account. Nodes with high eigenvector centrality are those that are connected to other nodes that, in turn, have high centrality. This measure captures the idea that influential nodes are those that are connected to other influential nodes, indicating a key role in facilitating information flow and network control^28^. Most previous analyses^28–30^ ignore negative correlations when calculating EC. However, in this study, we divided the dFC matrix into positive and negative matrices to prevent the loss of information. The EC of node *i* is defined as:

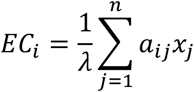

where *λ* is the eigenvalue of adjacency matrix.

All topological metrices were computed from three types of dFC matrixes: the original matrix, a matrix containing only positive dFC, and a matrix containing only negative dFC. This approach ensures the binary nature of the networks.

### 2.4 Statistical analysis

We compared the four summary statistics and dynamic topological metrics of dFC matrices between individuals with iRBD and HC, using the general linear model (GLM). The GLM defines a multivariate linear association between a set of explanatory/independent measures *X* and a set of outcome/dependent measures *Y*:

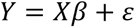

In the context of our MRI analyses, an outcome variable *Y* was dFC values recorded from the 132 VOIs of the 152 subjects. The explanatory variable *X* included the group (iRBD and HC), age and gender. We then evaluated effects of the group on the dFC or topological metrices.

### 2.5. Clinical scores and correlation analysis

We performed correlation analysis of statistically significant dFC metrics and the clinical batteries (including RBDSQ-J, MoCA-J and MDS-UPDRS part III). MoCA-J was available in all participants while the other batteries were available only in the iRBDs. For MoCA-J scores, we used GLM to associate them with age, gender, and disease. We use MoCA-J scores as Y, age, gender, and group (iRBD and HC) as X to evaluate the difference of MoCA-J score between groups and genders, also the relationship with age. Furthermore, we used the GLM to explore correlations between the statistically significant metrices and clinical assessments in the iRBD patients, when *Y* was the scores from RBDSQ-J, MoCA-J and MDS-UPDRS, and *X* included the age, gender, and a dFC or topological measure.

## 3. Results

### 3.1 Demographic data

Demographic information and behavioral test results are shown in Table 1. We accounted for the effect of gender difference between iRBD and HC groups, by adding gender as a covariate in all statistical tests.

**Table.1.** Participants’ geographic information. Data shown are the mean ±SD.

|  | Patients with RBD (n=55) | Healthy subjects (n=97) | Corrected p-value | T-value | Cohen's d |
| --- | --- | --- | --- | --- | --- |
| Age | 69.7±5.8 | 66.5±8.7 | 0.2515 | 2.0295 | 0.4588 |
| Sex(male/female)<sup>a</sup> | 39/16 | 47/50 | <0.001 | N.A. | N.A. |
| MoCA-J<sup>b</sup> | 24.9±3.0 | 24.8±3.2 | 0.4188 | -0.20382 | -0.01290 |
| MDS-UPDRS Part I | 4.6±3.4 | N.A. | N.A. | N.A. | N.A. |
| MDS-UPDRS Part II | 0.6±1.4 | N.A. | N.A. | N.A. | N.A. |
| MDS-UPDRS Part III | 1.8±2.0 | N.A. | N.A. | N.A. | N.A. |
| RBDSQ-J | 7.6±2.9 | N.A. | N.A. | N.A. | N.A. |

<sup>a</sup> Chi-square test was used to analysis the difference of gender proportion between iRBD and HC groups.
<sup>b</sup> MoCA-J scores of healthy subjects only involve 18 people.

**Table.2.** MoCA_J information.

|  | Corrected p-value | T-value | Cohen's d |
| --- | --- | --- | --- |
| Age | 0.2553 | -2.425 | N.A. |
| Sex | 0.3545 | N.A. | 0.5503 |
| Disease | 0.4188 | -0.20382 | -0.01290 |

### 3.2 Dynamic functional connectivity

Seventeen VOI pairs showed differences in mean dFC (Table 3a and Figure 1) and a VOI pair had a difference in kurtosis (Table 3b and Figure 2) (p < 0.05, FDR corrected). A decreased negative mean of dFC in iRBD (iRBD<HC) was found between the pre- or post-central gyrus and the posterior cingulate cortex or precuneus, precuneus and supplementary motor cortex, and supramarginal gyrus and middle temporal gyrus. Additionally, a weaker positive mean dFC in iRBDs (iRBD<HC) was observed within the cerebellum. Moreover, a stronger positive mean dFC in iRBDs (iRBD>HC) was shown between the cerebellum and pre- or post-central gyrus, thalamus, and anterior cingulate gyrus or Heschl’s gyrus, and amygdala and postcentral gyrus.

**Table.3.** Stronger/weaker mean (a) and kurtosis (b) of dFC in iRBDs compared to HCs. PC(posterior cingulate gyrus), PreCG r(precentral gyrus right), PreCG l(precentral gyrus left), PostCG r(postcentral gyrus right), PostCG l(precentral gyrus left), SMA r(supplementary moto area right), aMTG r(middle temporal gyrus, anterior division right), aSMG r (supramarginal gyrus, anterior division right), Thalamus r(thalamus right), AC(anterior cingulate gyrus), HG r (Heschl’s gyrus right), Ver8 (vermis 8), Cereb 45 r(cerebellum 45 right), Cereb 45 l(cerebellum 45 left), Cereb 6 r(cerebullum 6 right).

| Regions | Stronger/weaker dfc | Corrected p-value |
| --- | --- | --- |
| a. Mean of dFC |  |  |
| PC-PreCG r | RBD<HC | 0.04624 |
| PC-PostCG l | RBD<HC | 0.02626 |
| Precuneous- PreCG r | RBD<HC | 0.04775 |
| Precuneous-SMA r | RBD<HC | 0.03243 |
| aSMG r-aMTG r | RBD<HC | 0.04001 |
| Ver8-Cereb 45 r | RBD<HC | 0.03115 |
| Ver8-Cereb 45 l | RBD<HC | 0.03036 |
| Thalamus r-AC | RBD>HC | 0.03437 |
| Thalamus r-HG l | RBD>HC | 0.03065 |
| Amygdala l-PostCG r | RBD>HC | 0.03819 |
| Cereb 45 l-PreCG r | RBD>HC | 0.03205 |
| Cereb 45 l-PreCG l | RBD>HC | 0.03787 |
| Cereb 45 l-PostCG r | RBD>HC | 0.03226 |
| Cereb 45 l-PostCG l | RBD>HC | 0.01837 |
| Cereb 45 r-PreCG l | RBD>HC | 0.03389 |
| Cereb 45 r-PostCG l | RBD>HC | 0.0398006 |
| Cereb 6 r-PreCG l | RBD>HC | 0.03419 |
| b. Kurtosis of dFC |  |  |
| Cereb 9 r-aMTG r | RBD>HC | 0.04444 |

**Fig.1.**
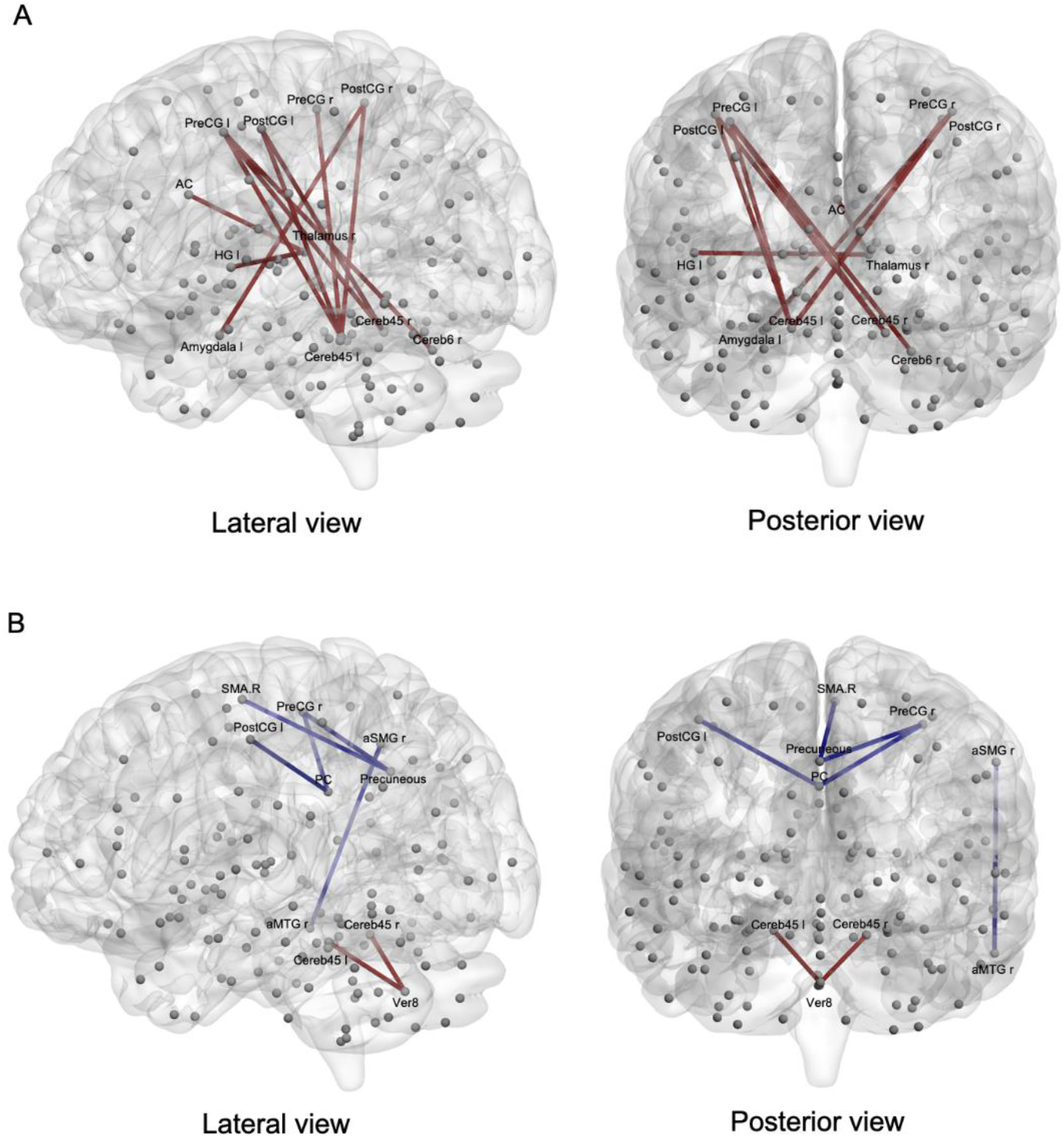
Differences of mean of dFC in iRBD and HC group (p<0.05, FDR corrected): (A) Stronger FC in iRBDs compared to HCs (iRBD>HC). (B) Weaker FC in iRBDs compared to HCs (iRBD<HC). Red(blue) lines in two panels indicating positive(negative) mean dFC in brain networks.

**Fig.2.**
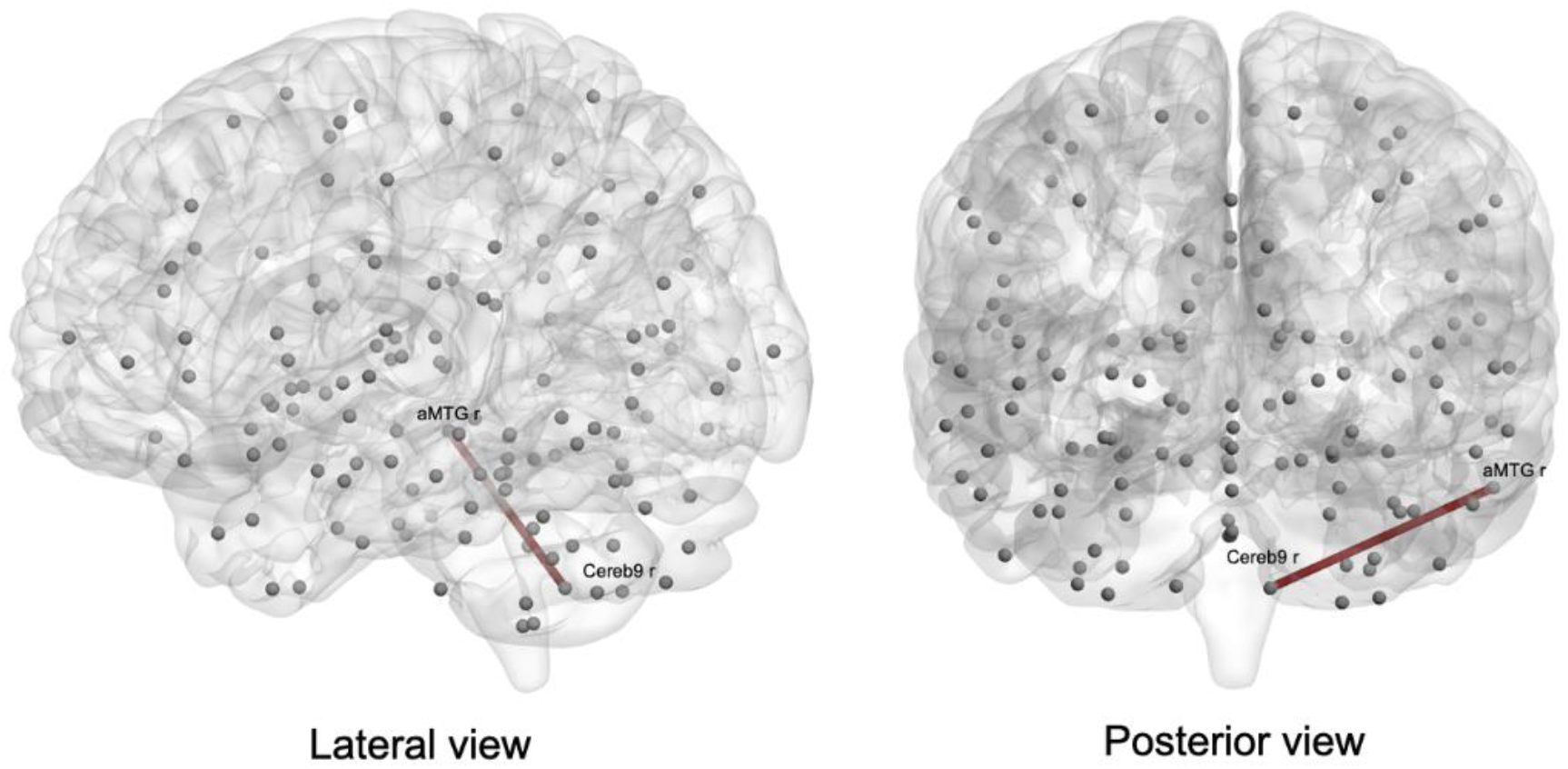
Difference of kurtosis of dFC in iRBD and HC group (p<0.05, FDR corrected). Red lines in two panels indicating stronger kurtosis FC in iRBD patients compared with HCs.

Furthermore, the kurtosis of dFC between the middle temporal gyrus and the cerebellum was higher in iRBDs than HC (iRBD>HC). The probability distribution of dFC values between the two regions of iRBDs and HCs were shown in figure 3. The left tail of the dFC distribution was thicker in individuals with iRBD compared to that of HCs, suggesting a greater occurrence of negative dFC values in iRBDs.

**Fig.3.**
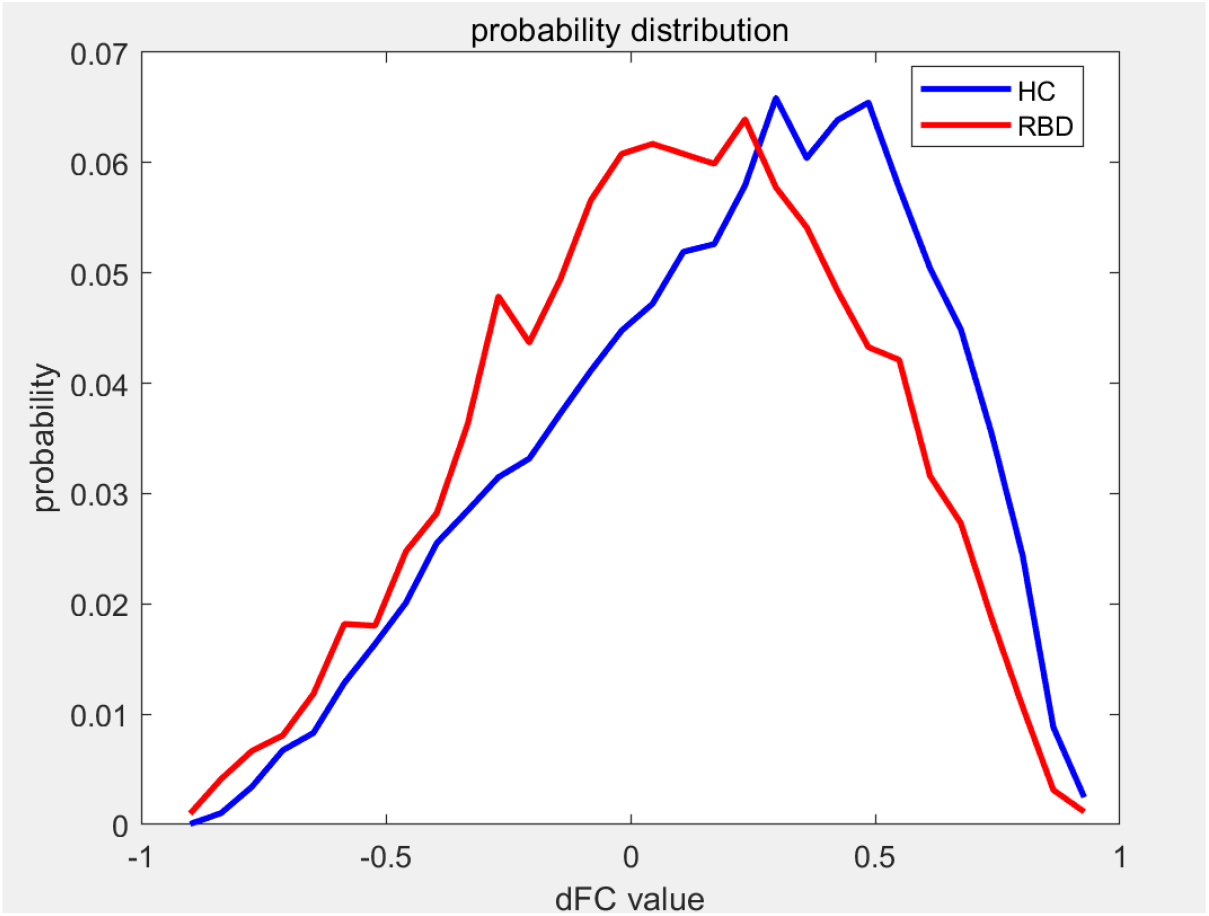
Probability distribution of dFC values. Red and blue lines indicating dFC values from iRBDs and HCs.

### 3.3. Graph theory analysis

Among the four topological metrics, only the result from the EC showed significant difference: the mean of eigenvector centrality of positive dFC matrix in iRBD patients increased in the right thalamus compared to HCs and the mean eigenvector centrality of negative dFC matrix in the iRBD decreased in the pre- and post-central gyri compared to HCs (Figure 4).

**Fig.4.**
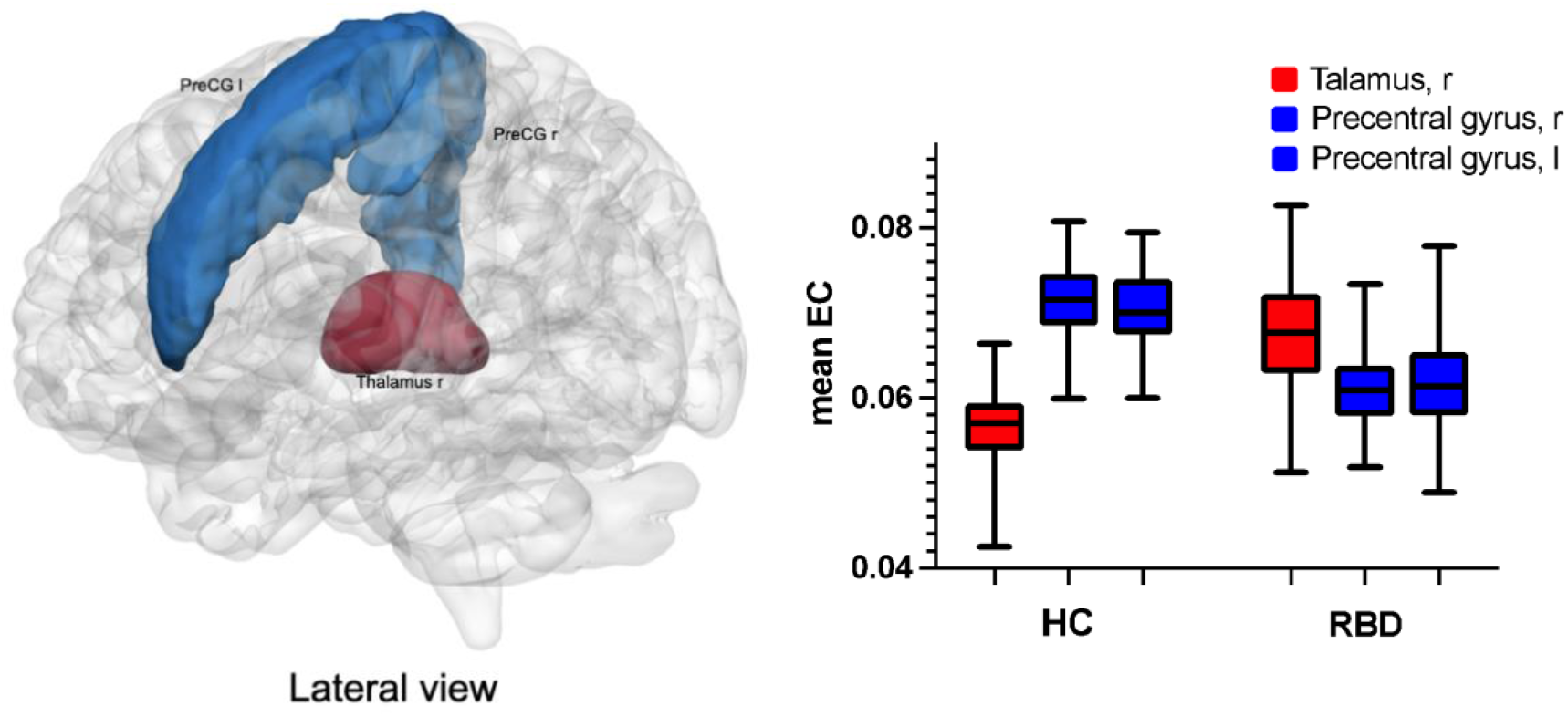
Differences of mean eigenvector centrality in iRBD and HC group (p<0.05, FDR corrected). Red/blue-colored regions indicating increased/decreased mean EC in iRBD patients compared with HCs.

### 3.4. Clinical scores and correlation

None of the 21 metrics above exhibited significant correlation with the clinical scores (RBDSQ-J, MoCA-J or MDS-UPDRS) after multiple comparisons (p<0.05, FDR corrected) (Table 2). The MoCA-J scores did not show significant relationship with disease, gender, or age, either (p<0.05, FDR corrected).

## 4. Discussion

Compared with HC, patients with iRBD showed altered mean dFC in the default mode network (DMN) including the posterior cingulate cortex and precuneus. Sensorimotor network (SMN), basal ganglia, and cerebellum also showed differences in mean dFC between the groups, replicating previous findings. The kurtosis of dFC between the middle temporal gyrus and the cerebellum showed group-wise differences, exemplifying novelty from our dFC approach. From the topological analysis, furthermore, the mean EC in the precentral gyrus and thalamus showed group-wise differences. These findings advanced our understanding of network-level pathophysiology of iRBD as prodromal alpha-synucleinopathy

### 4.1 Dynamic functional connectivity

#### 4.1.1. Mean dFC

The mean dFC analysis should basically echo findings from previous FC studies although differences in pre-processing and analytic methods may result in slight discordance across studies.

Compared to HC, we identified a weaker of negative mean dFC in iRBDs between parts of the DMN (including the precuneus/posterior cingulate gyrus) and the SMN, and this aberrant pattern implies disturbances in motor control and coordination. Analyses within the DMN suggest a link between the posterior cingulate cortex and the motor control circuits^31^, while the precuneus is associated with the coordination of motor behaviors^32^. Hence, we speculate that this decrease may indicate dysfunction in advanced movements controlled by the cognitive network (namely DMN) and possibly serve as a prodromal marker of PD, (and severe movement dysfunction is prevalent in developed PD). Abnormal mean dFC involving the SMN and DMN, as reported in previous FC studies in iRBD and PD^9,15,33,34,57^, is thus likely associated with latent motor and cognitive dysfunctions. It is plausible that the patients with iRBD were at a prodromal stage of movement disorders, implicating the possibility for developing further network abnormalities involving the SMN and DMN as the disease progresses.

HC showed positive mean dFC within the cerebellar network. We noted that such intra-cerebellar mean dFC was decreased in iRBD. By contrast, the positive mean dFC between the SMN and cerebellum was increased in iRBD compared with HC. Indeed, altered connectivity involving the cerebellum was shown as a key feature in our previous FC studies^9,57^. Weaker dFC of the intra-cerebellar network could underlie abnormal motor behaviors during sleep in iRBD, given a role of the cerebellum in regulating REM sleep^35^. Dysfunction in cerebellar connectivity has been linked to abnormal motor behaviors in iRBD^36^. The stronger dFC between the SMN and cerebellum may represent a compensatory mechanism, contributing to maintaining normal motor functions at the prodromal stage. Similarly stronger FC are rarely reported in iRBD^37^ than HC but are observed consistently in PD^38^. Our results replicate this stronger cerebellar-SMN FC in PD, highlighting the latent connection between iRBD and PD.

The connection between the thalamus and the anterior cingulate gyrus is integral to the multiple thalamo-cortical pathways and may play a crucial role in sensorimotor function and motor selection^39^. Additionally, the amygdala-motor pathway is believed to be involved in complex and subtle purposive behaviors^40^. Therefore, the stronger mean dFC between the thalamus and the anterior cingulate gyrus and between the amygdala and SMN in iRBD may be broaden the scope of compensation for motor functions. With regard to the amygdala, a structural MRI study has unveiled decreased volumes in the thalamus and amygdala in individuals with PD^41^. Given the close relationship of the amygdala to the development of anxiety and depression in the early stages of PD, the altered mean dFC related to the thalamus and amygdala may serve as a precursor to non-motor symptoms in PD-related disorders.

#### 4.1.2. Kurtosis of dFC

A novel finding was that the kurtosis of dFC between the middle temporal gyrus and the cerebellum was higher in individuals with iRBD than HCs. Kurtosis provides information not apparent in mean dFC. Kurtosis is a measure of “taildness” indicating how data distribute away from the normal distribution at ends, and higher kurtosis possibly indicates unusually dynamic changes in brain states^42^. For instance, kurtosis could be related to the instability of rs-fMRI signals^43^. The excess kurtosis in iRBD was primarily ascribed to unusually strong anti-correlations in iRBD (see Figure 4). Furthermore, the shape of the curve suggests that the distribution of HCs exhibits a prominent peak, while the curve representing iRBDs appears flatter and more dispersed, indicating a greater degree of dFC instability in iRBDs. According to past dynamic rs-fMRI analysis, kurtosis was believed to implicates a tendency for larger deviation^44^, In s previous rs-fMRI study, high kurtosis of dFC was reported in temporal lobe epilepsy, considered to show extreme dFC values related to cognitive dysfunctions^45^. Notably, an altered kurtosis of dFC between the middle temporal gyrus and cerebellum was reported in relation to abnormal cognitive functions in Alzheimer’s disease^46^. Put together, we hypothesize that the excess kurtosis indicates unusually strong anti-correlation and instability of dFC between the middle temporal gyrus and cerebellum in individuals with iRBD. Note that the mean dFC did not exhibit a difference between the middle temporal gyrus and cerebellum. Thus, changes in kurtosis may precedes the changes in the mean or occur independently from those. Considering the role of the temporal cortex in memory, excess kurtosis may serve a potential biomarker for early-stage cognitive dysfunction in iRBD from which a disease phenotype may covert to DLB that shares some aspects with Alzheimer’s disease. However, we need to exercise caution because this is the first application of kurtosis of dFC to rs-fMRI data from iRBDs and kurtosis has been criticized for its sensitivity to noise^43^. Therefore, further studies are needed to interpret the present results from kurtosis of dFC.

#### 4.1.3 Utility of dFC metrices

Here we employed summary statistics of dFC derived from time-series of rs-fMRI data after sliding window analysis, aiming to mitigate potential issues associated with the sliding window method and thereby obtaining robust findings. For dFC analysis, sliding window analysis is often followed by clustering or co-activation pattern analysis to explore various brain states through which resting-state brains may transit. Regarding iRBD, studies have highlighted differences between iRBDs and HCs in the temporal properties of these brain states^15,52^. However, recent studies have questioned the reliability of these analyses, by conducting similar analyses on simulated stationary data^43,53,54^. Note that these studies do not claim the non-existence of dynamics retrieved from sliding window analysis of dFC beyond static FC^54^. Although some popular dynamic features, such as state distribution and transition probability, can falsely emerge from simulated data without temporally varying data structure^53^, some nonlinear features including variance and graphical metrics cannot be explained with simulated data^55^. We thus avoided from using clustering analyses to define brain states; instead, we utilized summary statistics of non-Gaussian features of dFC time-series to interrogate if they have disease-related information.

Among the four moments of data distribution (i.e., mean, variance, skewness, and kurtosis), the mean dFC revealed the most robust differences between the groups as above. Conceptually, mean dFC is akin to static FC, suggesting that the results from the mean dFC should be similar to those from static FC analysis. Indeed, we replicated many findings from our previous static FC studies^9,57^. We however failed to find differences between the groups using variance and skewness measures. Actually, such a negative result aligns with the previous studies that showed static FC can explain most of the difference in connectivity between groups ^54,56^; yet, a small proportion of data variance cannot be extracted by static FC. Indeed, we found that kurtosis reflected a difference between iRBD and HC. Given its capability to measure non-Gaussian properties of dFC time-series and evidence from other diseases, kurtosis may serve as a promising feature for extracting genuine dynamic features reflecting network abnormality in neuro-psychiatric disorders.

### 4.2. Topological metrices

Another outcome of this study was the demonstration of increased mean EC of the thalamus coupled with decreased mean EC of the bilateral precentral gyri in iRBD compared to HC. To our knowledge, this study marks the first use of EC as topological measures to characterize network abnormality in iRBD. Excluding negative correlations in previous studies^28–30^ may lead to a loss of important information, as suggested by my results in the negative correlation network. EC estimates an importance of a node (i.e., VOI) while considering the whole-brain network, offering a more comprehensive understanding of the network structure beyond the ones possible with pairwise dFC/FC analysis. The decreased EC of the precentral gyri in the negative network, coupled with stronger mean dFC related to precentral gyrus, implies an altered pattern in the motor cortex: a loss of inhibition and a gain of activation, that maybe mark the early-stage motor function change in RBD. The thalamus serves not only as a relay station for information but also maintains extensive connections with the entire cerebral cortex, playing a crucial role in integrating information across multiple functional brain networks^47^. Given its role as a hub of the functional brain network, the heightened EC observed in the thalamus of individuals with iRBD may be interpreted again as a compensatory mechanism. This compensation potentially aims to uphold the overall brain network organization in iRBD, specifically by facilitating the global transfer and integration of information processing among cortical regions. Consistently, hyperactivation of the thalamus has been reported in iRBDs and PD^48,49^, which is accordacce with our results.

We did not observe group differences in clustering coefficients although altered clustering coefficients (C) in the precentral gyrus and thalamus have been reported in iRBDs and PD^5,51^. This may be due to the early stage of our iRBDs where local dysfunction might not be severe enough to be evident. We failed to find group-wise differences in global-level metrics including GE and assortativity coefficients. This may also be attributed to the early stage of iRBDs in our study where dysfunction is localized and well-compensated to maintain global performance. Topological analyses in previous iRBD studies have been limited, and the few available studies present contradictory results in GE^12,50^. This is according with our interpretation that different severity of iRBD may exhibit different global functions.

### 4.5. Limitations

Our study has some limitations. First, the heterogeneity in iRBD poses challenges in extracting reliable information. The limited results from the dynamic metrices may be an outcome of this heterogeneity when large difference in iRBDs reduced the signal detectability for possible biomarkers. This emphasizes the importance of follow-up studies of iRBD until final diagnosis. Furthermore, iRBD requires longitudinal and multimodal assessments in large samples to provide insights into progression and conversion to overt neurodegenerative diseases. Second, the clinical scores of HC might be insufficient to confirm the normal cognitive and behavioral functions. HCs only underwent the MMSE test, which is insensitive to subtle cognitive impairment. Therefore, we cannot affirm that HC were indeed cognitively intact. Third, our results may be biased because data were from one medical center, which may not be representative of the broader population,. Thirdly, I checked the small-worldness index of dFC matrixes and some of them did fit the small-worldness. So, the threshold used for the matrices may not be fully optimized for a graph theory analysis. Lastly, despite the observed results likely being related to motor or cognitive functional abnormalities, they did not correlate with the clinical scores including MDS-UPDRS or MoCA.

## Acknowledgments

I express profound appreciation to J-PPMI for collecting the data, a fundamental contribution to this study. I am also grateful for the participants who generously volunteered their time and cooperation, making this study possible. Their contribution is pivotal in advancing our understanding of neuroscience and neurodegenerative diseases. Furthermore, I extend my appreciation to Laboratory Integrated Neuroanatomy and Neuroimaging for providing the necessary resources and infrastructure essential for the successful execution of this research.

